# It takes extremes to be robust

**DOI:** 10.1101/478784

**Authors:** Felipe Bastos Rocha, Maria Dulcetti Vibranovski, Louis Bernard Klaczko

## Abstract

Phenotypic robustness is a central property of life, manifested in the ability of organisms to endure perturbing conditions throughout their development and often yield rather constant phenotypes. Fundamental questions on robustness (canalization) remain to be answered (see [1]). Do alleles that confer robustness against one perturbation also confer robustness to others? Is the robustness observed in multiple traits/taxa achieved through shared or specific mechanisms? Here, we describe an elementary model of trait development that yields phenotypic robustness without dedicated systems of developmental or transcriptional buffering. Robustness emerges when extremely low or high levels of gene activity lead to either depletion or saturation of the developmental system. We use this model to show that experimental results associating robustness to apparently redundant *cis-*regulatory sequences (from [2]) probably reflect a similar elementary system of saturation/depletion. We then analyze a large dataset of phenotypic responses of diverse traits of animals, plants and bacteria (from [3]) and show that the amount of response is mostly determined by the distance to the phenotypic extremes. Moreover, the most robust genotypes are often those that yield either extremely low or high phenotypes. Our results help reframing the concepts of canalization and plasticity, suggesting that phenotypic responses are mainly the result of variation in the very systems controlling each trait, rather than being attributable to either “plasticity genes” or “canalization genes”. Furthermore, they provide a hint on the causes of the genomic ubiquity of apparently redundant *cis*-regulatory sequences [4,5].

## Results and Discussion

Since Waddington’s seminal thesis on canalization [6], many authors assume that dedicated mechanisms that evolved to buffer development against perturbations are the major cause of the observed phenotypic robustness (e.g. [7,8]). Figure 1 (from *a* through *e*) illustrates how such mechanism may be represented in terms of a simple genotype-phenotype map where the phenotypic values are a linear function of gene activity. Perturbations of development are represented as forces that alter (either increasing or inhibiting) a genotype’s gene activity (Fig. 1b), and buffering mechanisms as any force that counteracts the potential effect of these perturbations (Fig. 1c). These two factors, combined with genetic variation for gene activity, yield a pattern of response variation where both responsive and robust genotypes may be observed at any value of the phenotypic axis (Fig. 1d and e).

**Figure 1.**
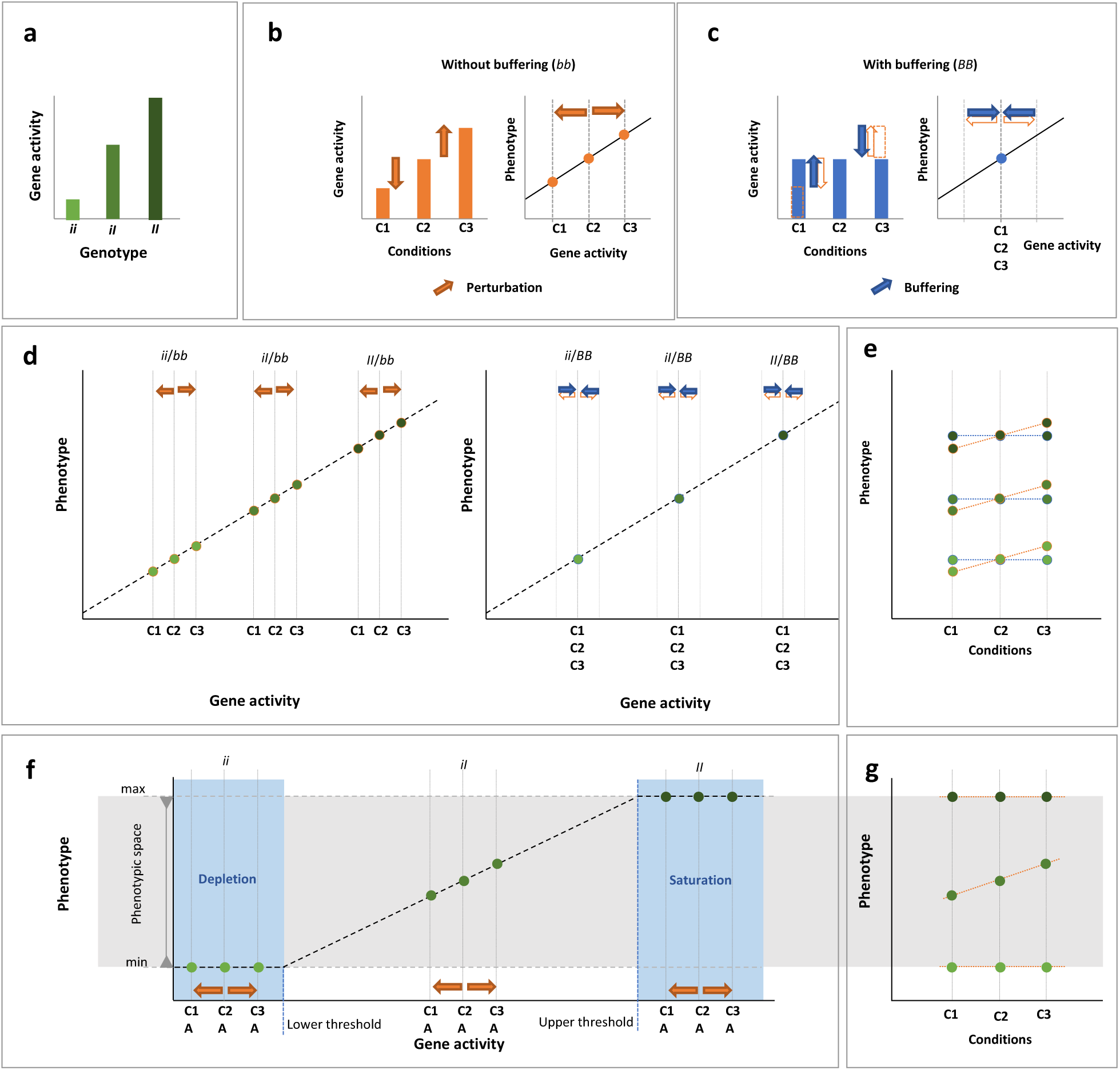
Robustness achieved by buffering and saturation/depletion dynamics yield contrasting patterns of variation of phenotypic responses to perturbations. **a.** three hypothetical genotypes (*ii, iI* and *II*) yield different mean levels of gene activity (green bars) for the gene *i*, which promotes the development of a continuous trait. **b.** three conditions – at the genetic background or the environment – (C1, C2 and C3) may affect the *i* gene activity (left) and consequently the phenotype of the trait it induces (right). **c.** this effect may be avoided by a second locus (*b*) that confers the ability to buffer against the effects of conditions C1 and C3 on gene activity (left) and thus stabilize the phenotype (right). **d.** the combined effects of the genetic variation for the inducer gene (*i*) and the buffering gene (*b*) shown in **a, b** and **c** yield contrasting patterns on a linear genotype-phenotype map. **e.** an independent buffering gene is capable of yielding stable response curves (blue dots) at any position of the phenotypic axis. **f.** a nonlinear genotype-phenotype map – here the simple consequence of a limited phenotypic space – yields robustness even when there is no buffering gene. The three genotypes shown, even though suffering the same amount of perturbation at the gene activity axis (orange arrows), but only the one with intermediary level of trait induction (*iI*) translates this perturbation at the phenotypic level. **g.** this creates a particular pattern of response variation where there is an association between the amount of phenotypic response and the position of the genotype on the phenotypic space.

Gene-activity/phenotype curves, however, are often nonlinear (Fig. 1f) [9,10] and thus yield nonlinear phenotypic responses when organisms are tested at several developmental conditions [11– 13]. Previous authors, from Plunkett [14] and Wright [15] to Hartl et al. [16] and Meiklejohn and Hartl [9] have already related the nonlinearity of dose-response curves in enzyme kinetics with instances of phenotypic robustness. While there is speculation on whether gene-activity/phenotype curves may evolve towards nonlinearity [1,10], this property may also be an intrinsic property of two general developmental features that operate at the extremes of the phenotypic space. First, for most (perhaps all) traits, the level of induction (here defined as the balance of activity between genes that promote and genes that inhibit a trait) must reach a threshold to yield phenotypic increments from the minimum possible phenotype. Second, all traits are only allowed to increase to a certain maximum phenotype. These properties create two regions in gene-activity/phenotype curves where changes in gene activity will not lead to changes in phenotype, because the developmental system is either depleted or saturated (Fig. 1f and g).

We built a simple model of trait development that explicitly incorporates these two developmental features and used it to assess how genotypes that confer different levels of gene activity respond to perturbations. Briefly, our model describes a trait formed by a limited group of cells that differentiate when the activity of a gene reaches or surpasses a fixed threshold. The phenotype is given by the number of differentiated cells. We modeled gene activity with two parameters: the probability of gene activation and the rate of transcription once the activation occurs [17,18]. We introduced genetic variation using variable basal levels of gene activity. We set perturbation as a function of a continuous environmental axis and used each genotype’s basal level of gene activity as the intercept of this function. We did not introduce any form of transcriptional or developmental buffering, i.e., all genotypes could potentially suffer the same effect from the environmental perturbation. We modeled gene activity following four setups. In setup (*1*), both environment and genetic variation affect the probability of gene activation (*A*); in setup (*2*), environment variation affects *A* and genetic variation affects the rate of transcription after activation (*T*); in setup (*3*) the environment affects *T* and genetic variation affects *A*; and in setup (*4*) both environment and genetic variation affect *T*.

We then evaluated the transcriptional and phenotypic reaction norms (Fig. 2) that would be produced by genotypes that were submitted to the same axis of environmental variation. We quantified the total amount of response in each reaction norm using Global Plasticity (GP) – the phenotypic range of each reaction norm (Fig. S1a) – to allow for diverse shapes and directions of response to be comparable (see [19] for further elaboration on this matter). Briefly, Global Plasticity allows to quantify the total amount of phenotypic change that a given set of conditions causes in genotypes irrespective of the direction, form or intensity of the response. When one is interested on the robustness of phenotypes, this is a better parameter than the traditional slope of reaction norms, for example, which carries the signal of the direction of the response and is not suitable for nonmonotonic responses. Using normalized values, genotypes with GP under 0.1 are those whose total response cover less than 1/10 of the maximum phenotypic range that is possible for that trait under the genetic system being analyzed. To assess the pattern of plasticity/robustness variation in each scenario, we plotted the transcriptional and phenotypic GP of genotypes against their respective indicators of mean gene activity: mean transcription across environments and mean phenotype across environments (see Methods).

**Figure 2.**
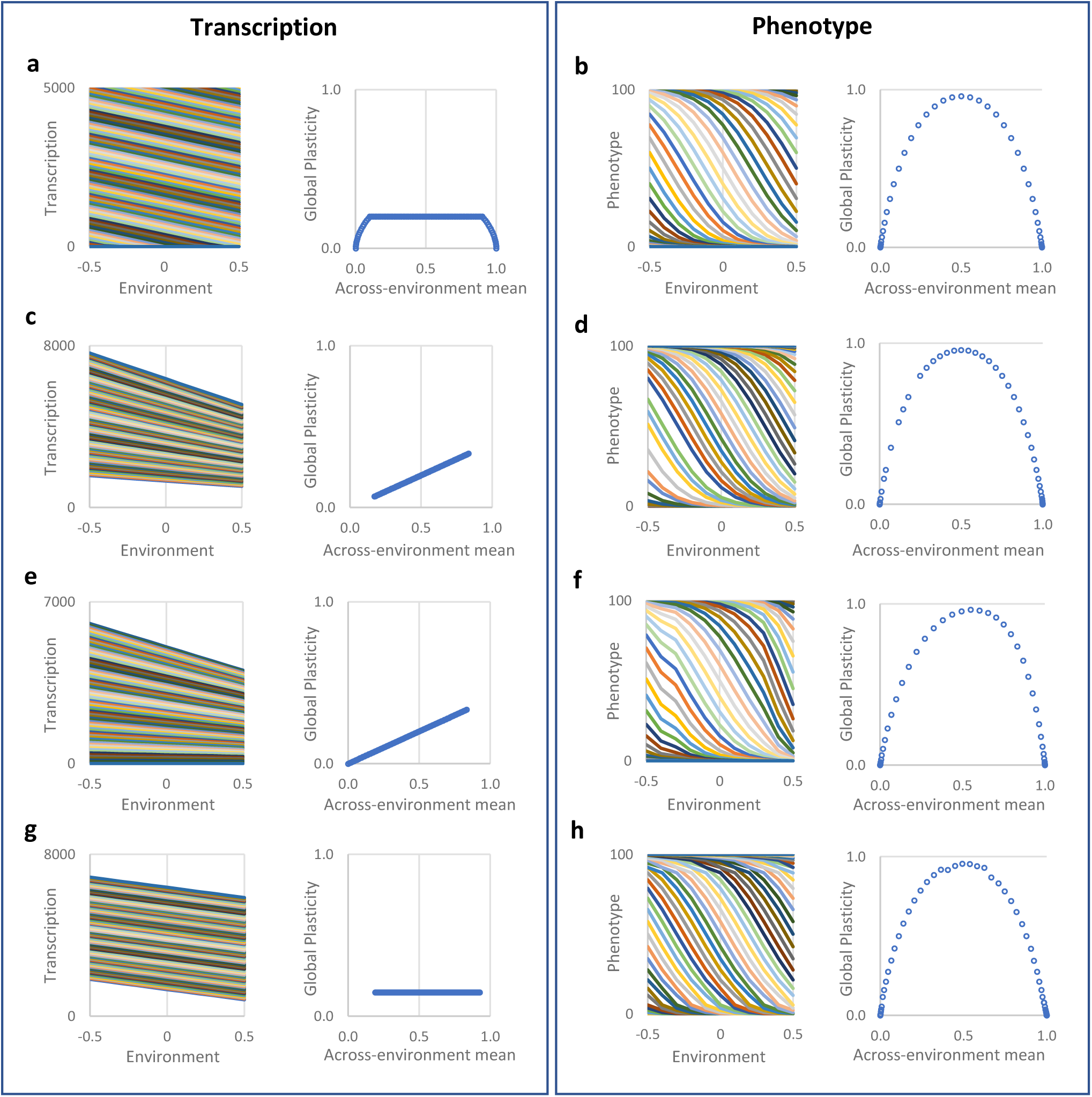
An elementary model of trait development suggests that gene activity and phenotype follow different rules when organisms are submitted to perturbations. Left column – transcriptional reaction norms and Global Plasticity by the across-environment mean transcription of each reaction norm. Right column – phenotypic reaction norms and Global Plasticity by the across-environment mean phenotype of each reaction norm. Results are grouped by gene activity setup: *i* (a and b) – environmental variation (Env.) and genetic variation (Gen.) affect the probability of gene activation (PGA); *ii* (c and d) – Env. affects PGA and Gen. affects the rate of transcription (RT); *iii* (e and f) Env. – RT, Gen. – PGA; *iv* (g and h) – Env. and Gen. – RT. Values shown correspond to *t* = 50 and *n* = 101. (See also Figures S1, S2 and S3).

The variations of transcriptional and phenotypic responses show a remarkable contrast (Fig 2 left and right halves, respectively). Transcriptional Global Plasticity is highly sensitive to the gene activity setup (Fig. 2, a,c,e and g). When both factors affect the Probability of Gene Activation (setup *1* - Env: PGA; Gen: PGA; Fig. 2a), transcriptional GP is constant for intermediary genotypes and decreases towards zero as the level of gene activity becomes either extremely low or high. In setups *2* (Env: PGA; Gen: RT; Fig. 2c) and *3* (Env: RT; Gen: PGA; Fig. 2e), the transcriptional GP linearly increases with the level of gene activity, yet with variable slopes. Finally, in setup *4* (Env: RT; Gen: RT; Fig. 2g) transcriptional GP becomes mostly independent on the genotype.

Variation of Phenotypic Global Plasticity, on the other hand, shows virtually the same pattern (although with slight variations) across all setups examined (Fig. 2b d, f and h): GP values gradually increase towards genotypes with intermediary mean phenotypes (i.e., those with intermediary gene activity) and decrease towards zero (i.e., become more robust) as the genotypes yield more extreme mean phenotypes. This relationship is highlighted if we convert the mean across-environment phenotypes into distance to the trait limits (DTL – see Fig. S3). This pattern holds if we use different values of environmental perturbation (Fig. S1) and different forms of perturbations (Fig. S2).

Therefore, in a system without a dedicated mechanism of developmental or transcriptional buffering, phenotypic stability is not a function of whether transcription is stable or how gene activity is controlled. It emerges as a consequence of properties of the developmental system that either becomes saturated or depleted due to extreme levels of gene activity, thus entailing a strong dependency on the distance to the trait limits (Fig S3). A genotype that is phenotypically robust due to saturation or depletion will therefore be robust to multiple forms of perturbations with comparable effects on gene activity: within-locus genetic perturbation (i.e., dominance), between-loci genetic perturbation, and multi-variate environmental perturbation (Fig. 1f and g), following Meiklejohn and Hartl’s [9] proposition that “*evolution should produce a single mode of canalization*”.

Is such a simple model pertinent to actual complex living organisms? Recent studies found that genotypes enriched for *cis*-regulatory sequences that drive similar patterns of expression (shadow enhancers [20–23] and homotypic clusters of transcription factor binding sites [2] tend to yield phenotypic stability before genetic and environmental perturbations. Crocker et al. [2], for instance, investigated the effect of cis-regulatory multiplicity on the robustness of larval trichomes of *Drosophila melanogaster*. They assessed how eight genotypes responded to variation at the genetic background (Ubx dose) and the environment (development at three temperatures). Each genotype had a specific arrangement of homotypic binding sites for the Ubx-Exd transcription factors in an enhancer driving expression of the gene shavenbaby (svb), with arrangements varying from zero to three functional binding sites. Their results show that the genotype with three unmutated binding sites is the one that keeps the highest level of expression when submitted to a lower Ubx dose and the one that keeps the highest phenotype across temperatures. These results resemble those of a previous study from Frankel et al. [21], who showed that having shadow enhancers controlling svb also yields robustness to genetic and environmental perturbation. Assuming that the phenotype is a reliable readout of gene activity, Crocker et al.[2] concluded that clusters of homotypic binding sites may confer regulatory robustness to the svb enhancer tested. This conclusion is in line with the hypothesis that multiple regulatory elements may provide gene activity robustness due to functional redundancy [24,25].

We pondered whether our model could improve the interpretation of Crocker et al.’ results. If cis-regulatory multiplicity (or the apparent redundancy) conferred regulatory robustness, we would expect a negative association between GP and the number of unmutated Ubx-Exd binding sites in the reaction norms described in their study. Using the graphical data from Crocker et al [2], we found that their results did not support this association; furthermore, they showed that the most robust genotypes had none or one Ubx-Exd unmutated binding site (Fig. 3a). Therefore, it could be the case that the number of binding sites had a cumulative effect on the level of induction of trichome formation, possibly due to increased activity of the *svb* gene. This would be consistent with experimental results showing that multiplicity of homotypic and heterotypic transcription factor binding sites and shadow enhancers may cumulatively increase their target gene activity [26–28]. Supporting this interpretation, the mean across-temperature phenotype was positively associated with the number of unmutated Ubx-Exd binding sites (Fig. 3b), thus raising the possibility that the variation of GP observed in Crocker et al.’s data was due to a system of saturation/depletion.

**Figure 3.**
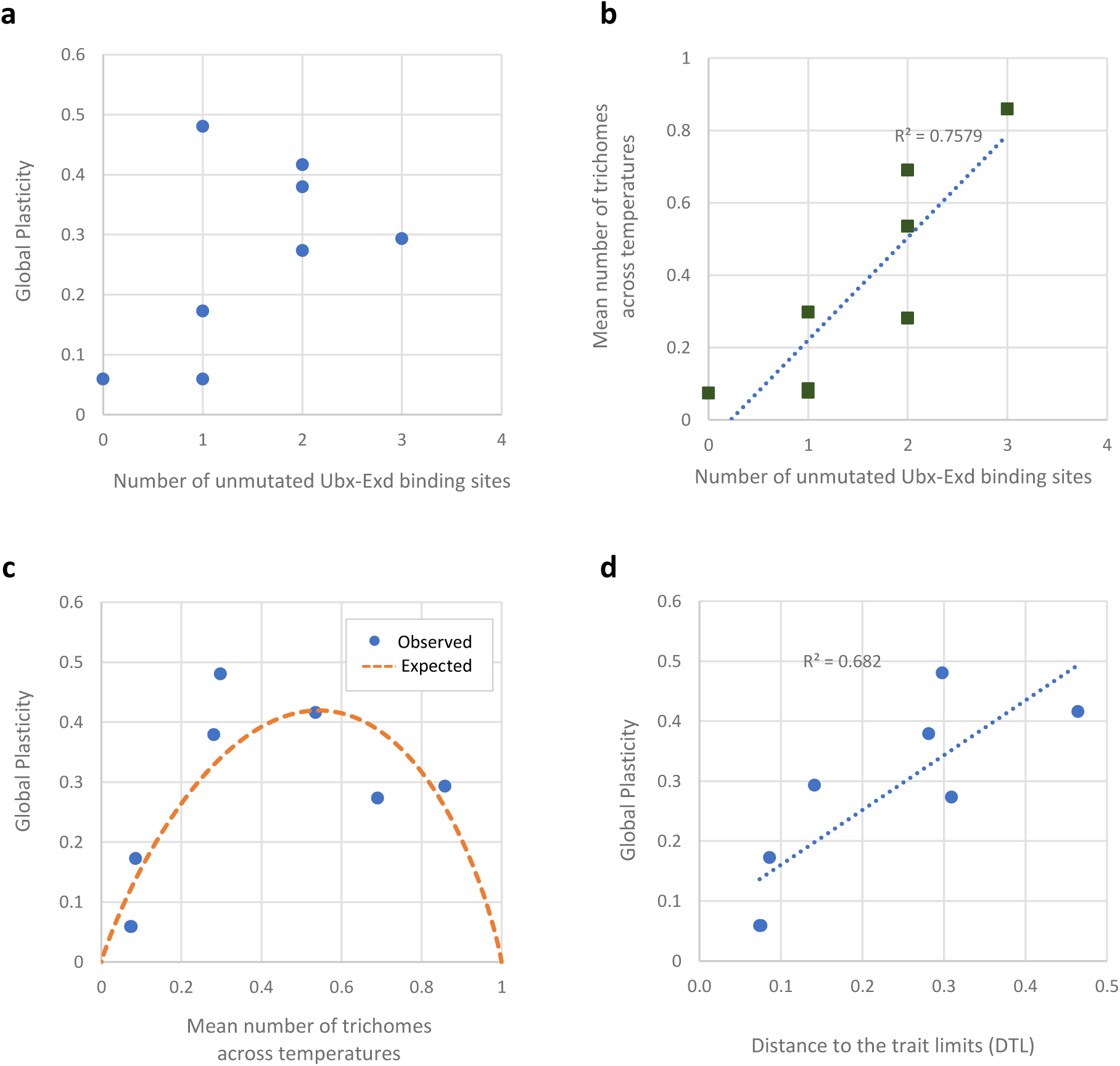
The association between *cis*-regulatory multiplicity and robustness described by Crocker et al. (2015) is consistent with a model of robustness by saturation and depletion. **a** – the lack of a pattern of variation of Global Plasticity by the number of Ubx-Exd binding sites does not support a positive association between cis-regulatory multiplicity and robustness. **b –** the mean number of larval trichomes is positively associated with the number of binding sites. **c –** observed values of Global Plasticity values are well predicted by a model of saturation depletion, and are significantly associated with the distance to the trait limits (DTL – **d**).

To evaluate this possibility, we plotted the GP of the eight RNs described by Crocker et al.[2] against each genotype’s across-temperature mean phenotype (Fig. 3c blue dots), which showed a bell-shape very similar to the expected by our model. Accordingly, GP variation could be significantly explained by a positive linear function of the distance to the trait limits (Fig. 3d). We used our model to estimate the expected GP values assuming that svb activity was perturbed by temperature at the same intensity in all genotypes. Surprisingly, even under this very simplistic assumption, our model offered a good prediction of the observed GP values (R^2^ = 0.679 – Fig. 3c).

These results offer a more nuanced view on how phenotypic robustness may be produced by *cis*-regulatory multiplicity, where robustness is achieved via a two-step process. First, a *cis*-regulatory sequence enriched for homotypic clusters of binding sites and/or shadow enhancers maximizes its target gene’s activity. As the target gene activity increases and eventually surpasses the saturation threshold at the genotype-phenotype curve, the genotype becomes phenotypically robust to any sort of perturbation that reduces the target gene activity without crossing back the saturation point (Fig. 1f). In terms of the system analyzed by Crocker et al. (2015) and previously by Frankel et al. [21], this means that both shadow enhancers and the homotypic binding sites within at least one of them may act in the same manner, incrementing the basal level of *svb* activity above the saturation point and thus protecting the phenotype from both environmental and genetic perturbations.

Interestingly, if our model indeed describes processes underlying the association between cis-regulatory sequences and phenotypic robustness, we expect the signal of this association to vary depending on the range of values of trait induction analyzed. At least two studies seem to follow this prediction. Preger-Ben Noon et al. [29] found that a robust low phenotype in the larval trichome system was achieved through the loss of shadow enhancers and gain of repressor binding sites in *D. sechellia*. At the genomic level, Paixão and Azevedo [30] found a negative relationship between cis-regulatory multiplicity and robustness in yeast, possibly reflecting a predominance of robustness by depletion over robustness by saturation in this organism.

Even if these results reflect the biological principles outlined in our model, does a system where phenotypic robustness is only achieved by saturation or depletion provide any clue on the mechanisms underlying the observed genetic variation of plasticity or canalization across virtually all organisms? We could expect systems where GP variation is dominated by saturation/depletion to show a mix of curves of GP variation as a function of the distance to the trait limits (e.g. Fig S3). Following this rationale, we analyzed 648 reaction norms from 34 animal species (76 traits), 15 plant species (36 traits) and one bacteria (two traits) from a dataset compiled by Murren et al. [3]. To make these diverse responses comparable, we normalized the reaction norms by each species-specific trait, and all environmental values by experiment, so that all reaction norms fitted a 1x×1 plot (see Methods for details). The general pattern of GP variation strongly resembled that of our model (Fig. 4a). Accordingly, GP values were highly dependent on the distance to the trait limits (R^2^ = 0.541; F = 762.861; p < 0.0001 -Fig. 4b). This association was significant across experimental designs and organisms (Sup Fig. 2), suggesting a key contribution of saturation/depletion (~ 50%) for the variation of response captured by the studies surveyed.

**Figure 4.**
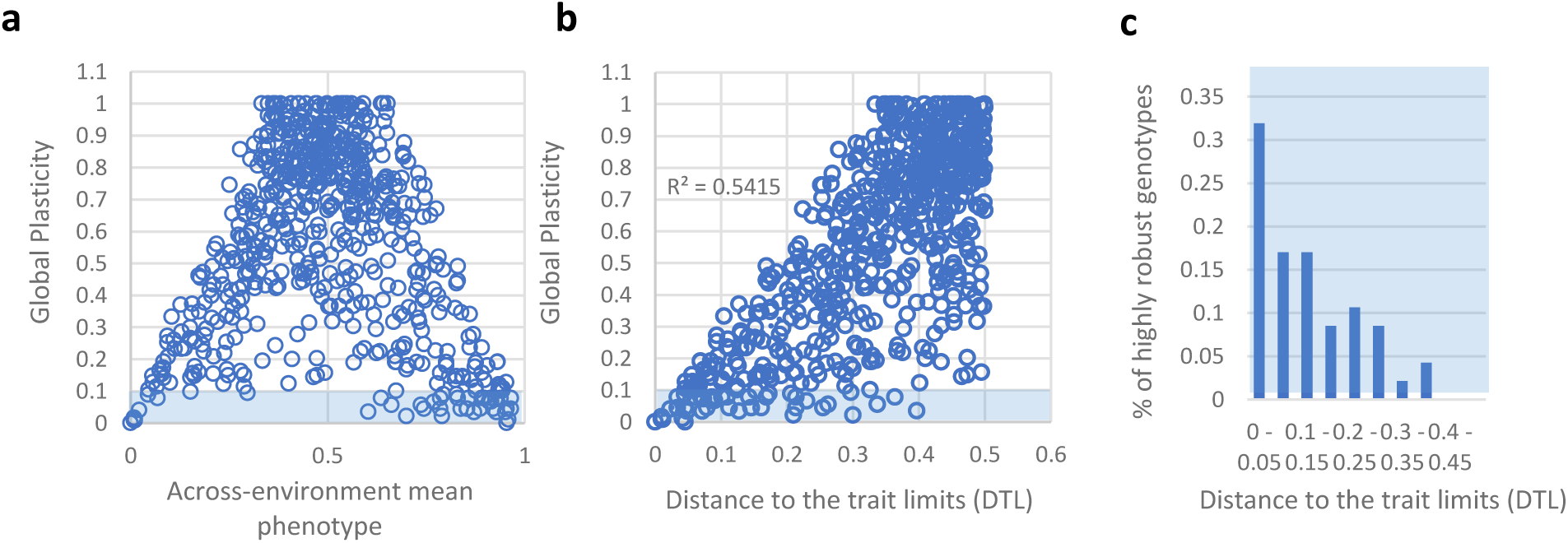
Phenotypic robustness variation in a sample of 648 reaction norms of diverse animal, plant and bacteria species is mostly explained by the distance to the trait limits. **a**, observed Global Plascitiy increases towards intermediary mean phenotypic values and decreases towards zero towards extreme phenotypes in the dataset extracted from Murren et al. (2014). **b,** this results in a significant dependency on the distance between the genotype’s mean phenotype and the trait limits. **c**, highly robust genotypes (those with Global Plasticity up to 0.1) are significantly concentrated near the phenotypic extremes in this sample of reaction norms. (See also Figure S4).

As expected, GP values disperse as they increase, which could mean that part of GP variation is due to possible buffering mechanisms consistent with the paradigmatic concept of canalization. Still, such dispersal can also be due to variation among species-specific traits, i.e., the fact that not all traits will be able to yield a GP of 1 even in the absence of saturation or depletion (see Figs. S1, S2 and S3 for an illustration of this effect). It is reasonable to assume that GP dispersal in such a myriad of traits and species must be due to a combination of between-trait variation (i.e., variation without buffering) and within-trait variation (i.e., variation due to buffering). In any case, extremely robust genotypes, as characterized by reaction norms with GP equal or lower than 0.1, were strongly concentrated towards extreme phenotypes (more than 74% of these genotypes with DTL ≤ 0.2), significantly deviating from a uniform distribution (Fig. 4c – D = 0.310, p < 0.0001). These results are substantial evidence that saturation/depletion dynamics is the leading cause of phenotypic robustness in this dataset.

The fact that such a clear pattern emerges from such a diverse set of developmental systems implies first developmental principles as the likely cause of both trait and response variation. Interestingly, both shadow enhancers and clusters of homotypic binding sites seem to be widespread across species genomes [4,5]. Whether this ubiquity is always related to phenotypic robustness is still unclear. Future investigation on this issue could profit from taking into account the nonmonotonic relationship between gene activity and robustness that is entailed by depletion/saturation dynamics.

It is worth noting that the generality of the recent findings on cis-regulatory variation and the reframing of concepts proposed here may have important implications for the study of complex diseases. G. Gibson [31] proposed that complex diseases may emerge as a result of decanalization, which led to the proposition that robustness should be considered in GWAS designs, possibly reducing the “missing heritability” [32]. Among the several implications of our findings to this matter, we may cite the change from an organism-level concept of robustness to a trait-level concept and the predictability introduced by saturation/depletion dynamics into the concept of genotype-environment interaction.

The model outlined here drives a change on the view of how much can organisms control their response to environmental perturbations. Our findings suggest that they are more often than not unable to buffer the developmental processes underlying phenotype against perturbations. This is in marked contrast with the predominant view on the evolution of phenotypic responses, that often assumes organisms to be masters of their development, equipped with mechanisms that modulate how and how much their phenotype will change. This conflict of views has a direct relationship with the genetic basis of phenotypic responses. From the plasticity perspective, many authors assume that the genetic basis of plasticity is independent of the genetic basis of the trait itself [33–35]. From the canalization/robustness perspective, the same assumption is often held, especially since the discovery of the buffering abilities of the chaperone Hsp90 [36–38]. Our results, on the other hand, imply that, for a given trait in a given species, the locus of response variation (call it plasticity or robustness/canalization) is more likely to control the variation of the trait itself than a separate system of response control (cf. [19,39]). The predominant role of cis-regulatory evolution underlying phenotypic change is a key general principle of evo-devo [40,41]. Accordingly, cis-regulatory changes are more capable of modulating the target gene activity with specificity of time and space than other forms of control of gene activity [42]. The consistency between our model and results regarding *cis*-regulatory architecture in various systems suggests that this may be a cardinal rule of the phenotypic robustness evolution.

## Acknowledgments

We thank the financial support of the following: Coordenação de Aperfeiçoamento de Pessoal de Nível Superior (CAPES), Conselho Nacional de Desenvolvimento Científico e Tecnológico (CNPq) and Fundação de Amparo à Pesquisa do Estado de São Paulo (FAPESP). FBR was supported by a postdoctoral fellowship from FAPESP (processo n° 2013/04980-0) and a fellowship from the Programa Nacional de Pós-Doutorado (PNPD) from CAPES. MDV is supported by a Young Investigator Award (processo n° 2015/20844-4). FBR thanks S. Scheiner, F. Uno and I.M. Ventura for comments on the manuscript and F. Uno for the title suggestion.

## Declaration of Interests

The authors declare no conflict of interest.

## Supplemental figures

**Figure S1.**
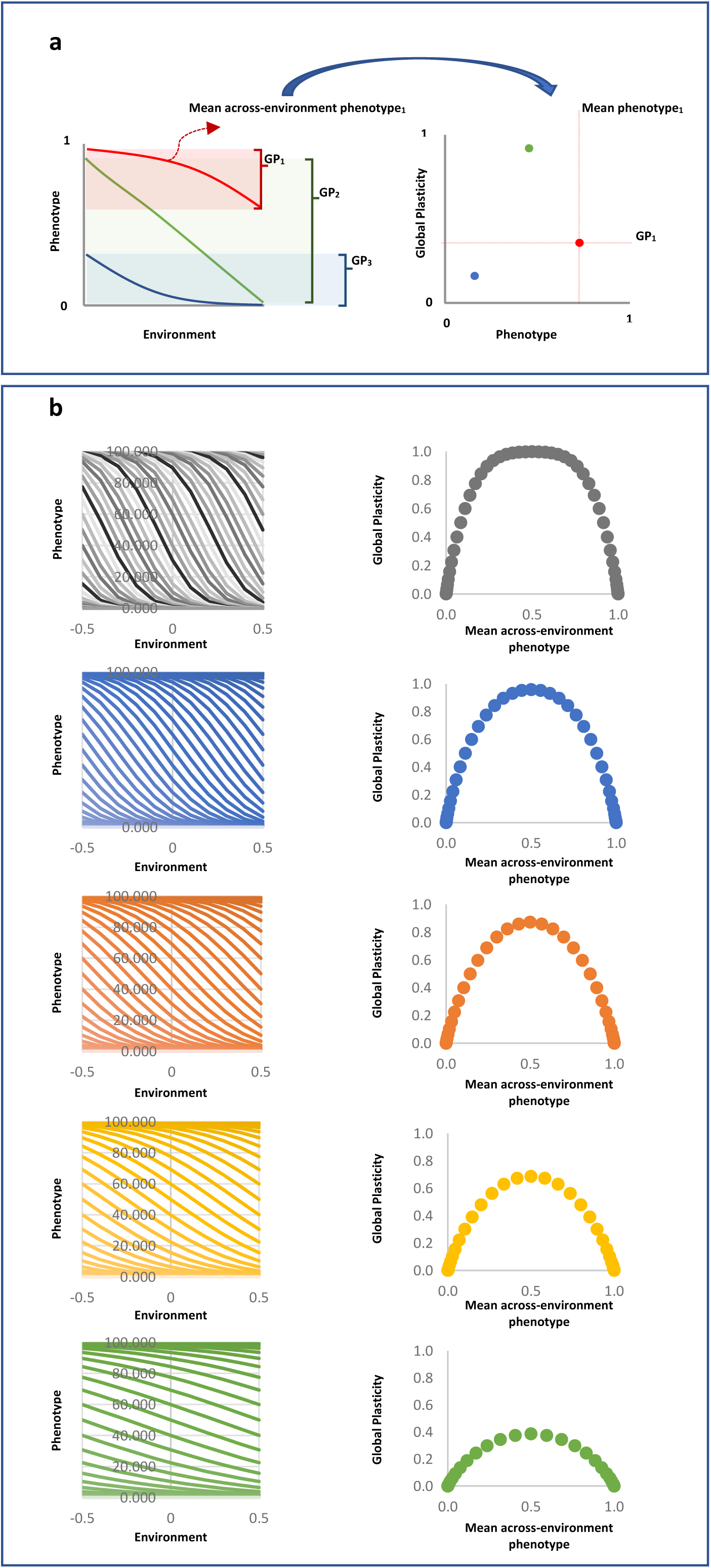
Phenotypic reaction norms produced by diverse values of environmental perturbation yield different patterns of Global Plasticity variation across the phenotypic space. **a –** illustration of the relationship between reaction norms and the pattern of Global Plasticity variation. **b –** phenotypic reaction norms yielded by decreasing values of environmental perturbation (left column) and their respective patterns of Global Plasticity variation.

**Figure S2.**
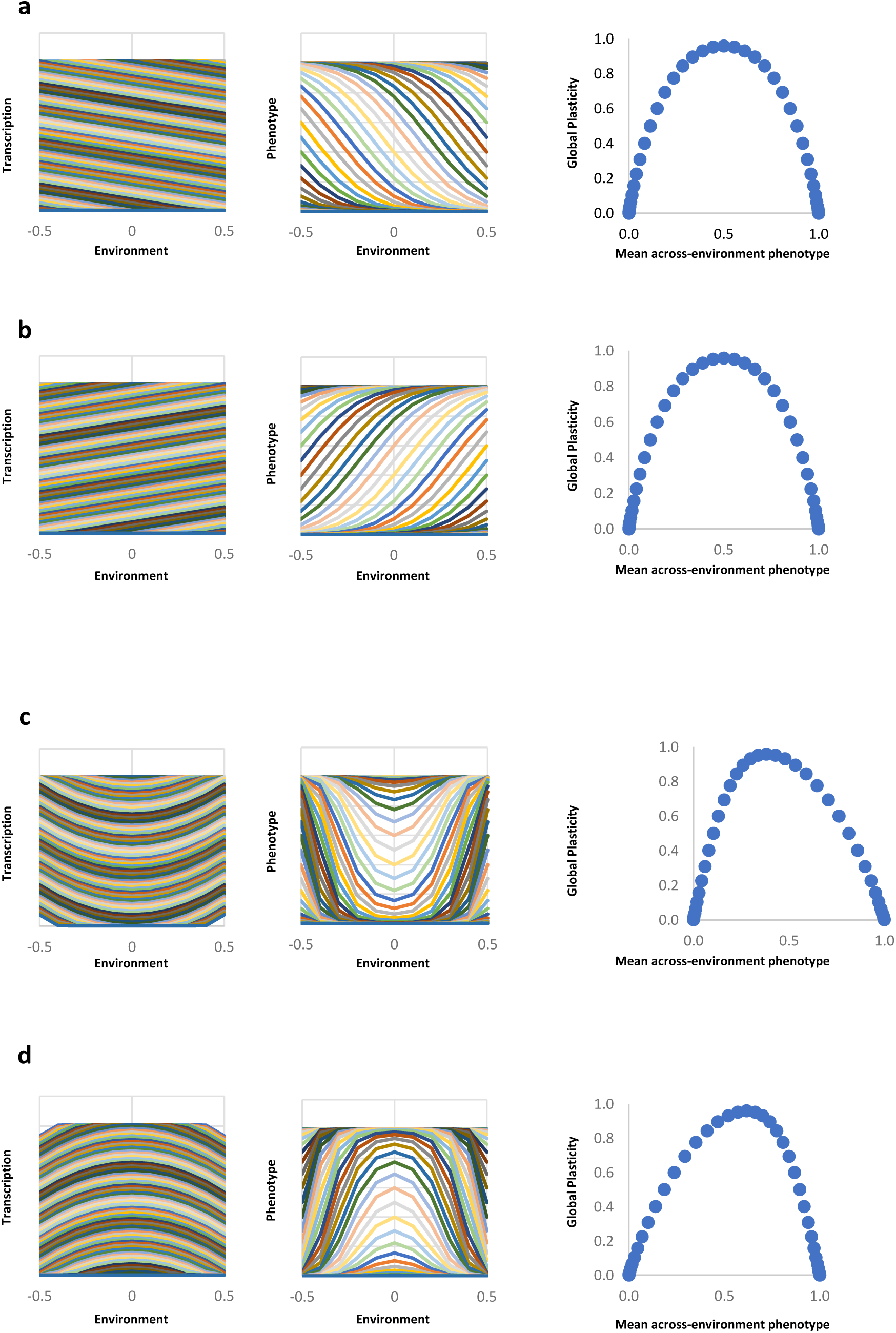
Different forms of environmental perturbation yield rather similar patterns of Global Plasticity variation. **a** – negative linear perturbation. **b** – positive linear perturbation. **c** – positive quadratic perturbation. **d** – negative quadratic perturbation.

**Figure S3.**
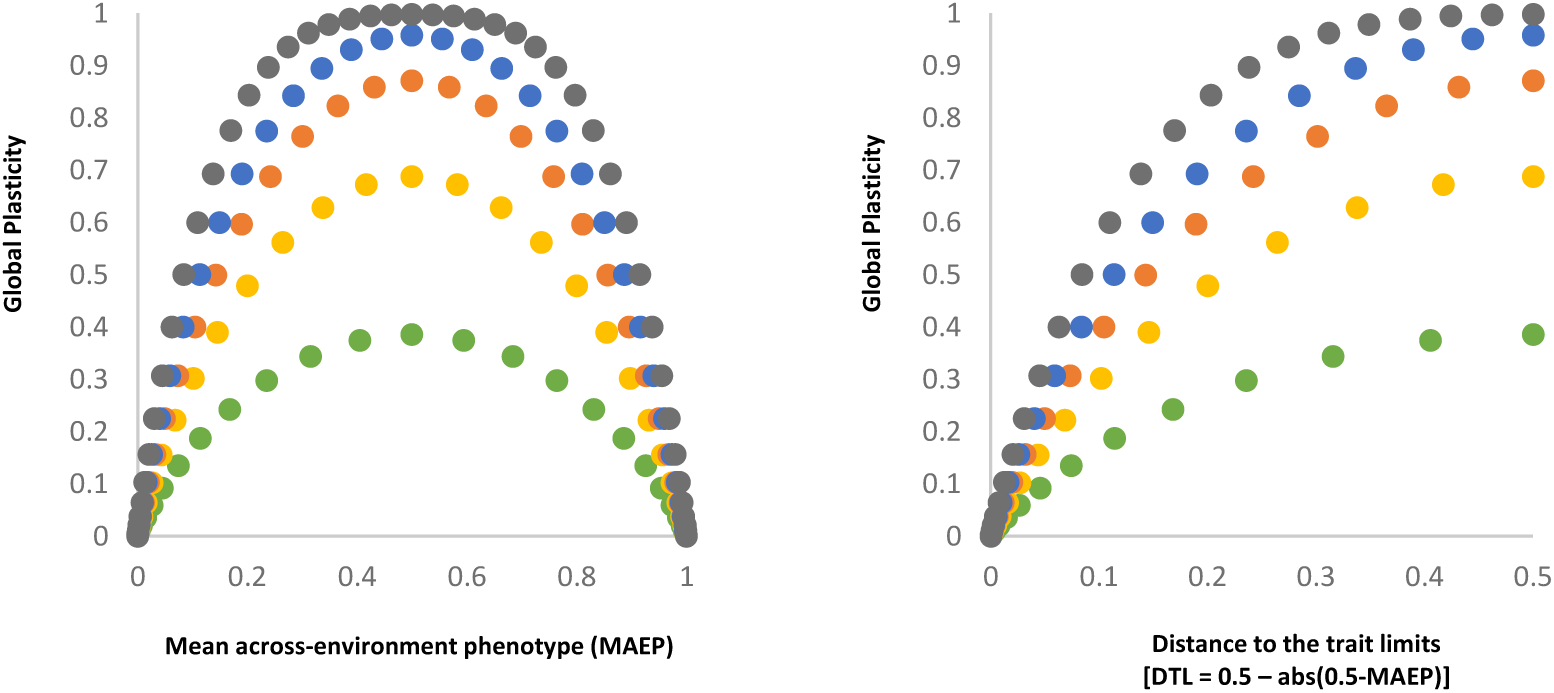
The superimposition of the patterns of Global Plasticity variation with different environmental perturbation values illustrates the effect of variation (whether among traits or genotypes) on the variation as a function of the mean across-environment phenotype (left) or the distance to the trait limits (right).

**Figure S4.**
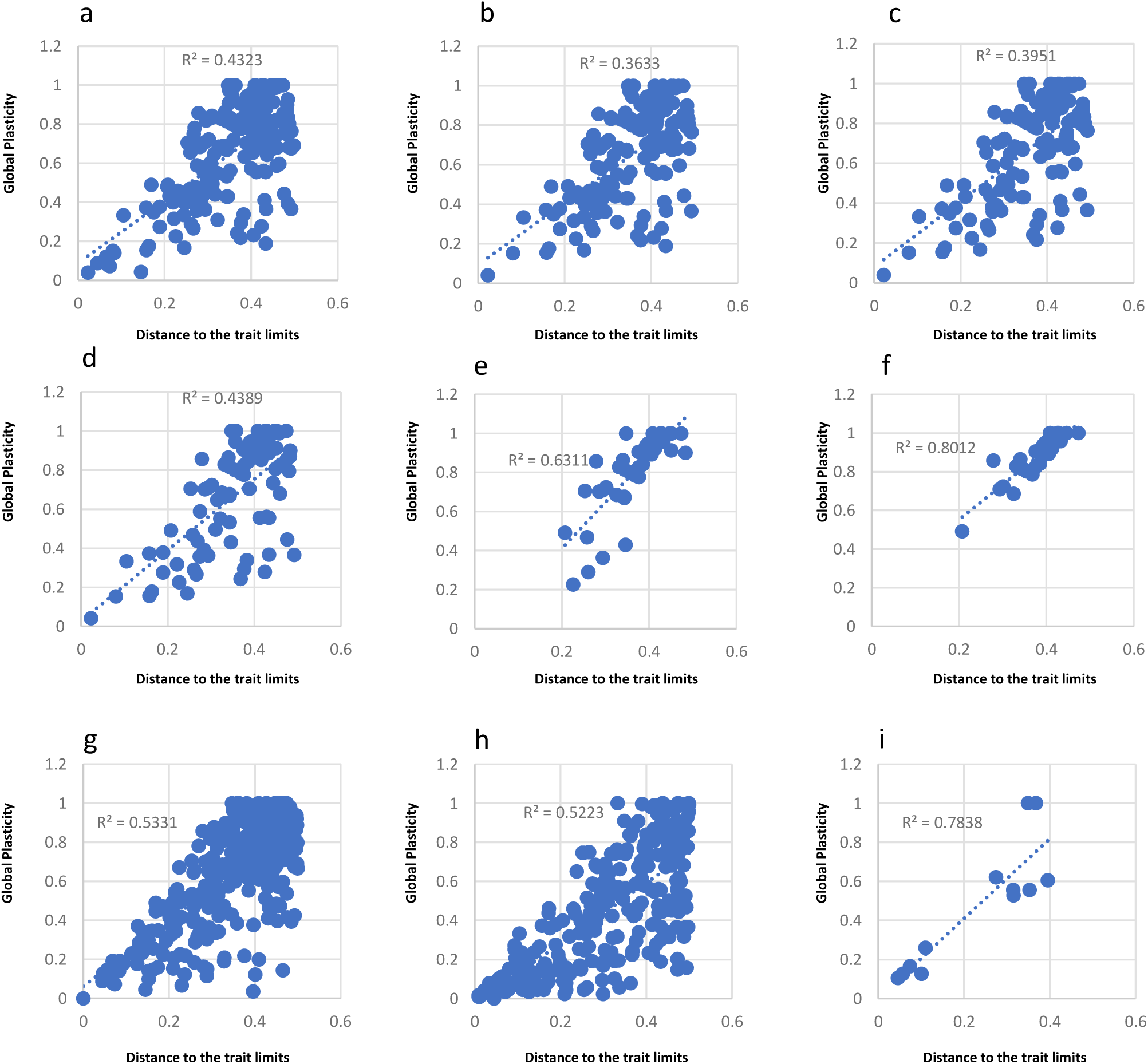
The pattern of association between Global Plasticity variation from Murren et al. 2014 (Fig.4) is consistent across different experimental setups and taxa. **a** trough **f** – Global Plasticity variation of reaction norms described with minimum number of environmental conditions varying from four through nine, respectively. **g, h** and **i** – Global Plasticity variation of animal, plant and bacteria species, respectively.

## Methods

### Model

The model describes the mean phenotype given by the number of cells following a path of differentiation that is conditioned by the probability of reaching a fixed threshold (*t*) of activity of an Inducer Gene (IG) in an infinite population. Each individual has a constant number of 100 cells that are capable of expressing IG, representing a constant trans-regulatory landscape of cells that constitutively express the transcription factor necessary for the IG activity. There is a developmental time-window during which the IG expression may trigger differentiation. There is a fixed maximum number of times (*n*) each cell may or may not activate the transcription of the IG (rounds of activation) during this developmental time window. Whether transcription is activated at each round of activation during the developmental time-window depends on the probability of activation (*p*). Thus, the probability of each cell to activate IG *k* times (*Pr*) is given by the binomial distribution:

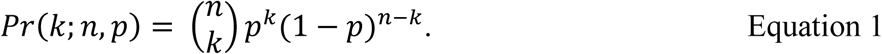

The realized transcription (*RT*) in each cell is given by the product between the number of times that IG is activated (*k*) within the developmental time-window and the rate of transcription (*r*). Hence, the phenotypic value of an individual is given by the number of cells activating the IG at or above the minimum number of activations necessary to reach *t*, or *k*_min_, which is given by

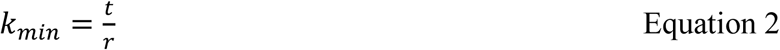

Genetic variation was introduced by varying uniformly either the probability of transcription initiation at the environment 0 (*A*) or the rate of transcription at the environment 0 (*T)*. The environmental effect was introduced by setting either *p* or *r* to be linear functions of a continuous range of environments (*E*) varying between −0.5 and 0.5 (given a linear perturbation, the results of GP for positive and negative functions are the same). Therefore, *p* and *r* were modeled as:

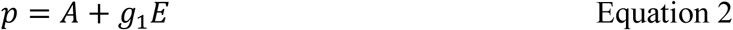

and

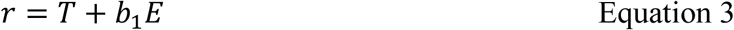

We evaluated the impact of four setups of gene activity by varying the parameters of these two equations. In setup *1,* multiple values of *A* were evaluated, *T* was fixed, and *g*_*1*_ was non-zero and *b*_*1*_ was zero. In setup *2, A* was fixed, *T* varied, *g*_*1*_ was non-zero and *b*_*1*_ was zero. In setup *3*, multiple values of *A* were evaluated, *T* was fixed, *g*_*1*_ was zero and *b*_*1*_ was non-zero. Finally, setup *4* used *A* fixed, *T* varying, *g*_*1*_ was zero and *b*_*1*_ was non-zero.

We considered that all cells of all individuals of a given genotype (i.e., a given value of *T* and *A*) that develop at a given environment have the same *p* and *r*. Therefore, the proportion of cells activating IG *k* times in an infinite number of individuals raised in each environmental value is given by the binomial probability *Pr*(*k*;*n,p*).

Accordingly, transcriptional reaction norms were evaluated by calculating the mean Realized Transcription 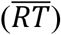 as the product of the mean number of activations (*p* · *n*) by the rate of transcription (*r*):

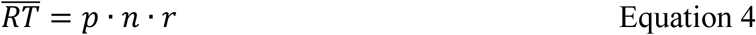

Likewise, the mean phenotype in each environment may be calculated from the product of the proportion of cells activating IG at or above *k*_min_ by the total number of cells. We calculated this from the complement of the cumulative probability of activating IG up to *k*_min_ −1 times:

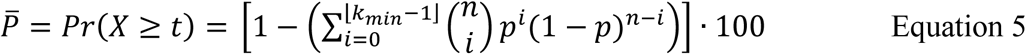

Different developmental setups, varying *t, n* and the total number of cells produced the same qualitative results. The results shown correspond to *t* = 2500 and *n* = 101. Likewise, we tested various values of *g*_*1*_ and *b*_*1*_ and assessed the impact of a second-degree environmental perturbation. While some of these changes do alter the shape of transcriptional and phenotypic reaction norms, they do not alter the overall shape of the curve of Phenotypic Global Plasticity variation (Fig. S2).

Transcriptional and phenotypic reaction norms were drawn as the mean RT and mean P for each genotype (i.e., each combination of A and T), respectively in each environment from −0.5 to 0.5, with 0.1 increments. For each reaction norm, transcriptional and phenotypic GP were calculated as the difference between the maximum and the minimum values of RT and P. The mean across-environment values of RT and P were the simple average of all values of each reaction norm. To facilitate the visualization of the relationship between phenotypic GP and the mean across-environments phenotype (MAEP), we converted MAEP values into the mean distance from the trait limits (DTL) as:

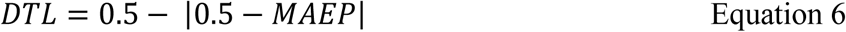

### Adjustment to the dataset extracted from Crocker et al. 2015

We captured each reaction norm point available in Figure 6 from Crocker et al. [2] using tps.dig [43] and converted the values to normalized phenotypic and environmental values. We calculated the GP and MAEP for each reaction norm. We tested the association between MAEP and the number of unmutated Ubx-Exd binding sites in each genotype, as well as between GP and the number of unmutated binding sites. We also performed a linear regression of the GP values on the *DTL* to test the dependency of plasticity on the distance to the limits of the trait.

In our model, each specific slope of perturbation (either *g*_*1*_ or *b*_*1*_) generated a curve of GP variation as a function of MAEP with a specific height and shape. These curves converge at extreme phenotypes (due to saturation and depletion) and diverge at intermediary mean phenotypes, where the peak of GP values occur (Fig. 3).

We used the setup *i* of our model to estimate the expected GP from the MAEP of Crocker et al.’s [2] reaction norms. Since their curves were mainly nonmonotonic with a peak phenotype at the intermediary temperature, we set *p* to be a negative quadratic function of the environment:

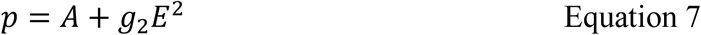

In this configuration, we verified that the relationship between *g*_*2*_ and GP could be well described by a second order polynomial (*g*_*2*_ = −0.3982 + 1.1111 GP - 1.636 GP^2^; R^2^ = 0.9918). We used this relationship to calibrate our model, estimating the *g*_*2*_ that would be necessary to yield the GP of the genotype whose MAEP was closest to 0.5 (. We used the model calibrated with this *g*_*2*_ (−0.21916) to estimate the expected GP values of the remaining genotypes (i.e., those not involved in the model calibration) based on their mean phenotype only and calculated the *R*^2^ between observed and predicted values to evaluate the goodness of fit of our model to their experimental results.

### Analysis of reaction norms from the dataset compiled by Murren et al. [3]

We retrieved the data compiled by Murren et al. from Dryad [44] and filtered for reaction norms that had continuous (non-qualitative) environmental variables and more than two reaction norms by trait in each species. This preliminary assessment resulted in a dataset of 648 reaction norms. We then normalized all environmental values by experiment and all phenotypic values by trait and species so that all reaction norms were fitted within a 1×1 plot. For each species-specific trait, the upper and lower extreme values were estimated by the maximum and minimum observed phenotypes regardless of genotype or environment. GP, MAEP and DTL were calculated for each reaction norm as described above. We tested the dependency of GP on DTL by performing a least-squares linear regression. To assess whether the regression was influenced by the number the fact that many experiments had only tested three environments, we repeated this analysis using only experiments with progressively higher minimum number of environments. All regressions were positive and significant. We tested the distribution of highly robust genotypes (i.e., the reaction norms with GP lower or equal to 0.1) against a uniform distribution using a Kolmogorov-Smirnov test.

